# Overexpression of sedoheptulose-1,7-bisphosphatase enhances photosynthesis in *Chlamydomonas reinhardtii* and has no effect on the abundance of other Calvin-Benson cycle enzymes

**DOI:** 10.1101/2020.02.14.949040

**Authors:** Alexander Hammel, Frederik Sommer, David Zimmer, Mark Stitt, Timo Mühlhaus, Michael Schroda

## Abstract

The productivity of plants and microalgae needs to be increased to feed the growing world population and to promote the development of a low-carbon economy. This goal can be achieved by improving photosynthesis via genetic engineering. In this study, we have employed the Modular Cloning strategy to overexpress the Calvin-Benson cycle (CBC) enzyme sedoheptulose-1,7 bisphosphatase (SBP1) up to 3-fold in the unicellular green alga *Chlamydomonas reinhardtii*. The protein derived from the nuclear transgene represented ∼0.3% of total cell protein. Photosynthetic rate and growth were significantly increased in SBP1-overexpressing lines under high-light and high-CO_2_ conditions. Absolute quantification of the abundance of all other CBC enzymes by the QconCAT approach revealed no consistent differences between SBP1-overexpressing lines and the recipient strain. This analysis also revealed that the eleven CBC enzymes represent 11.9% of total cell protein in *Chlamydomonas*. Here the range of concentrations of CBC enzymes turned out to be much larger than estimated earlier, with a 128-fold difference between the most abundant CBC protein (rbcL) and the least abundant (triose phosphate isomerase). Accordingly, the concentrations of the CBC intermediates are often but not always higher than the binding site concentrations of the enzymes for which they act as substrates. The enzymes with highest substrate to binding site ratios might represent good candidates for overexpression in subsequent engineering steps.

## 1 Introduction

An increased productivity of plants and microalgae is required to feed the growing world population and to promote the development of a low-carbon economy. One way to increase plant and microalgal productivity is to improve photosynthesis by genetic engineering. Engineering efforts that have resulted in increased biomass are the rewiring of photorespiration (Kebeish et al., 2007;Nolke et al., 2014), the improvement of linear electron transport between the photosystems (Chida et al., 2007;Simkin et al., 2017b), or the overexpression of distinct Calvin-Benson-Cycle (CBC) enzymes (for recent reviews see Simkin et al. (2019) and Kubis and Bar-Even (2019)). The rationale behind the latter approach is that the rising concentration of atmospheric CO_2_ caused by the burning of fossil fuels increases the velocity of the carboxylation reaction of Rubisco and inhibits the competing oxygenation reaction. This results in a shift in the limitation of photosynthesis away from carboxylation of ribulose 1,5-bisphosphate (RuBP) and towards RuBP regeneration. The CBC enzyme sedoheptulose-1,7 bisphosphatase (SBPase) has been shown to exert strong metabolic control over RuBP regeneration at light saturation, as it is positioned at the branch point between regenerative (RuBP regeneration) and assimilatory (starch biosynthesis) portions of the CBC. SBPase catalyzes the irreversible dephosphorylation of sedoheptulose1,7-bisphosphate (SBP) to sedoheptulose-7-phosphate (S7P). Accordingly, the overexpression of SBPase alone (Lefebvre et al., 2005;Tamoi et al., 2006;Feng et al., 2007;Rosenthal et al., 2011;Fang et al., 2012;Ding et al., 2016;Driever et al., 2017;Simkin et al., 2017a) or of the cyanobacterial bifunctional SBPase/FBPase (BiBPase) (Miyagawa et al., 2001;Yabuta et al., 2008;Ichikawa et al., 2010;Gong et al., 2015;Ogawa et al., 2015;Kohler et al., 2017;De Porcellinis et al., 2018) resulted in marked increases in photosynthesis and biomass yields in tobacco, lettuce, *Arabidopsis thaliana*, wheat, tomato, rice, soybean, and in the microalgae *Synechococcus, Euglena gracilis* and *Dunaliella bardawil*.

Genetic engineering often is an iterative process essentially consisting of four steps: (i) the design and manufacturing of a gene construct, (ii) its transfection into the target organism and the recovery of transgenic lines, (iii) the screening for expressing transformants, and (iv) the readout of the trait to be altered, on which basis the gene construct for the next cycle is designed. The cloning steps used to be a time constraint, which was overcome by new cloning strategies like Gibson assembly or Modular Cloning (MoClo) for Synthetic Biology that allow the directed assembly of multiple genetic parts in a single reaction (Gibson et al., 2009;Weber et al., 2011). Still a major time constraint (months) in the genetic engineering of plants is the recovery of a transfected plant and its propagation for reading out the altered trait. This constraint can only be overcome by using plant models with short generation times, like microalgae.

A potential problem of genetic engineering are undesired side effects of the genetic engineering that can best be revealed by system-wide approaches. One way is to compare the proteomes of wild type and engineered lines by quantitative proteomics (Gillet et al., 2016). A more targeted approach is the use of so-called quantification concatamers (QconCATs) (Beynon et al., 2005;Pratt et al., 2006). QconCATs consist of concatenated proteotypic peptides, an affinity tag allowing purification under denaturing conditions and, optionally, amino acids like cysteine or tryptophan for easy quantification. The QconCAT protein is expressed with a heavy label in *E. coli* from an *in silico* designed, codon-optimized synthetic gene cloned into an expression vector. A known amount of the QconCAT protein is then added to the sample and, upon tryptic digestion, the heavy proteotypic peptides from the QconCAT protein are released together with the corresponding light peptides from the parent proteins. All QconCAT peptides are present in a strict 1:1 ratio at the concentration determined for the entire protein. After ionization, the pairs of heavy QconCAT peptides and light native peptides can be separated and quantified by mass spectrometry, with the heavy peptides serving as calibrators allowing absolute quantification of the target proteins in the sample. This method is limited to about 20 targets per QconCAT protein.

The aim of this work was to provide a proof of principle for a rapid metabolic engineering workflow to improve photosynthesis. We chose to overexpress SBPase via the MoClo strategy, the unicellular green alga *Chlamydomonas* as a chassis, and QconCAT-based absolute quantification as a tool for monitoring effects on other CBC enzymes.

## 2 Material and Methods

### 2.1 Growth of *Chlamydomonas* cells

*Chlamydomonas reinhardtii* UVM4 cells (Neupert et al., 2009) were grown in Tris-Acetate-Phosphate (TAP) medium (Kropat et al., 2011) on a rotatory shaker. For transformation, cells were grown at a light intensity of 100 μmol photons m^−2^ s^−1^ to a density of 5 × 10^6^ cells/ml and collected by centrifugation at 4000 g for 2 min. 5 × 10^7^ cells were mixed with 1 µg DNA linearized with *Not*I and transformed by vortexing with glass beads (Kindle, 1990). Vortexed cells were diluted 2-fold with TAP and 2.5 × 10^7^ cells were spread onto TAP agar plates containing 100 μg ml^−1^ spectinomycin. Plates were incubated over-night in the dark and then incubated at 30 μmol photons m^−2^ s^−1^ for about ten days. For growth curves, cells were inoculated in 100 ml TAP medium and grown at 150 µmol photons m^−2^ s^−1^ to a density of about 8 × 10^6^ cells/ml. 100 ml TAP or Hepes-Minimal-Phosphate (HMP) medium (5 mM Hepes-KOH instead of 20 mM Tris, no acetate) were then inoculated with 3 × 10^5^ cells/ml in triplicates for each strain and growth was monitored by cell counting using the Z2 Coulter Particle Count and Size Analyzer (Beckmann). The culture volume is the summed cell volume of all cells in one ml medium. For mass spectrometry analyses, samples were harvested 22 h after inoculation (early log phase).

### 2.2 Measurement of oxygen evolution

Cells were inoculated in 50 ml TAP medium and grown overnight to early log phase. Oxygen measurements were performed in the Mini-PAM-II (Walz, Germany) device using the needle-type oxygen microsensor OXR-50 (Pyroscience, Germany). Before the measurements, the cell density was determined, and an aliquot was taken to determine the chlorophyll concentration. The PAM chamber was filled with 400 µl of *Chlamydomonas* culture and NaHCO_3_ was added to a final concentration of 30 mM. Cells were dark-adapted for 5 min and far-red light adapted for another 5 min. Then light with the intensities of 16, 29, 42, 58, 80, 122, 183, 269, 400, 525, 741 and 963 µmol photons m^−2^ s^−1^ was applied for 2 min each and oxygen evolution was monitored.

### 2.3 Cloning of the *Chlamydomonas SBP1* gene for MoClo

Our constructs are based on the Phytozome 12 annotation of the genomic version of the *Chlamydomonas SBP1* gene (Cre03.g185550) with seven exons and six introns. However, we used the first ATG in the 5’ UTR as start codon instead of the third proposed by Phytozome. To domesticate a BsaI recognition site in the fifth exon (GAGACC → GAGACA), the *SBP1* gene was PCR-amplified on total *Chlamydomonas* DNA in two fragments with primers 5’-TT**GAAGAC**ATAATGGCCGCTATGATGATGC-3’ and 5’-AC**GAAGAC**GGGTTGTCTCCTTGACGTGC-3’ for fragment 1 (1257 bp) and with primers 5’-TT**GAAGAC**GGCAACCCACATCGGTGAG-3’ and 5’-TT**GAAGAC**TCCGAACCGGCAGCCACCTTCTCAGAG-3’ for fragment 2 (963 bp; BpiI sites in bold letters). PCR was done with Q5 High-Fidelity Polymerase (NEB) following the manufacturer’s instructions and in the presence of 10% DMSO. The two PCR products were combined with destination vector pAGM1287 (Weber et al., 2011), digested with BpiI and directionally assembled by ligation into level 0 construct pMBS516. The latter was then combined with plasmids pCM0-020 (*HSP70A-RBCS2* promoter + 5’UTR), pCM0-101 (MultiStop) or pCM0-100 (3xHA), and pCM0-119 (*RPL23* 3’UTR) from the *Chlamydomonas* MoClo kit (Crozet et al., 2018) as well as with destination vector pICH47742 (Weber et al., 2011), digested with BsaI and ligated to generate level 1 constructs pMBS517 (L1-SBP1-mStop) and pMBS518 (L1-SBP1-3xHA). Both level 1 constructs were then combined with pCM1-01 (level 1 construct with the *aadA* gene conferring resistance to spectinomycin flanked by the *PSAD* promoter and terminator) from the *Chlamydomonas* MoClo kit, with plasmid pICH41744 containing the proper end-linker, and with destination vector pAGM4673 (Weber et al., 2011), digested with BpiI, and ligated to yield level 2 constructs pMBS519 (*aadA*+*SBP1*-mStop) and pMBS520 (*aadA*+*SBP1*-3xHA). Correct cloning was verified by Sanger sequencing

### 2.4 Screening of SBP1 overexpressing lines

Transformants were grown in TAP medium until mid-log phase and harvested by centrifugation at 13,000 g for 5 min at 25°C. Cells were resuspended in DTT-carbonate buffer (100 mM DTT; 100 mM Na_2_CO_3_), supplemented with SDS and sucrose at final concentrations of 2 % and 12 %, respectively, vortexed, heated to 95°C for 5 min, and centrifuged at 13,000 g for 5 min at 25°C. The chlorophyll content was determined as described by (Vernon, 1960). Total proteins according to 1.5 µg total *Chlamydomonas* chlorophyll were loaded on a 12 % SDS-polyacrylamide gel and analyzed by immunoblotting using a mouse anti-HA antibody (Sigma H9658, 1:10,000) for transformants with SBP1-3xHA or a rabbit anti-SBPase antibody (Agrisera AS15 2873, 1:2,500) for SBP1-mStop. Detection was done via enhanced chemiluminescence using the FUSION-FX7 device (Peqlab).

### 2.5 QconCAT protein expression and purification

The coding sequence for the Calvin-Benson cycle QconCAT protein (CBC-Qprot) was codon-optimized for *E. coli*, synthesized by Biocat (Heidelberg) harboring BamHI/HindIII restriction sites, cloned into the pET-21b expression vector (Novagen), and transformed into *E. coli* ER2566 cells (New England Biolabs). Expression of CBC-Qprot as a ^15^N-labeled protein and purification via Co-NTA affinity chromatography and electroelution was performed as described previously for the photosynthesis QconCAT protein (PS-Qprot) (Hammel et al., 2018). The eluted protein was concentrated, and the buffer changed to 6 M urea using Amicon ultra-15 centrifugal filter units with 10,000 MWCO (Merck). The protein concentration was determined spectroscopically at 280 nm on a NanoDrop™ spectrophotometer based on the Lambert-Beer’s law assuming a molecular weight of the CBC-Qprot of 47921 Da and an extinction coefficient of 37,400 M^−1^ cm^−1^. The protein concentration was adjusted to 1 μg/μl and the protein was stored at −20°C.

### 2.6 In solution tryptic digest and LC-MS/MS analysis

Twenty micrograms of total *Chlamydomonas* protein, as determined by the Lowry assay (Lowry et al., 1951), were mixed with 12.5, 25, 50, and 100 ng CBC- and PS-Qprot for replicates 1-3, and with 25, 50, 100, and 200 ng CBC- and PS-Qprot for replicates 4-6. Proteins were then precipitated with ice-cold 80% acetone overnight and digested as described previously (Hammel et al., 2018). 1 μg of the CBC-Qprot was precipitated with acetone without *Chlamydomonas* protein to obtain ion chromatograms for the Q-peptides alone. Tryptic peptides corresponding to 10 µg were desalted on home-made C18-STAGE tips (Empore) as described by (Rappsilber et al., 2007), eluted with 80% acetonitrile/2% formic acid, dried to completion in a speed vac and stored at −20°C. Peptides were resuspended in a solution of 2% acetonitrile, 2% formic acid just before the LC-MS/MS run. The LC-MS/MS system (Eksigent nanoLC 425 coupled to a TripleTOF 6600, ABSciex) was operated in μ-flow mode using a 25 μ-emitter needle in the ESI source. Peptides were separated by reversed phase (Triart C18, 5 μm particles, 0.5 mm × 5 mm as trapping column and Triart C18, 3 μm particles, 300 μm × 150 mm as analytical column, YMC) using a flow rate of 4 μl/min and gradients from 2 to 35% HPLC buffer B (buffer A: 2% acetonitrile, 0.1% formic acid; buffer B: 90% acetonitrile, 0.1% formic acid). The efficiency of ^15^N incorporation in the labeled peptides was estimated according to (Schaff et al., 2008). The intensities for the monoisotopic, fully ^15^N labeled peak (Mi) and the preceding, first unlabeled peak (Mi-1), containing one ^14^N, were extracted using PeakView v2.2 software (ABSciex) and used for calculating the labeling efficiency (99.39 ± 0.37% (SD)). BioFsharp was used for the extraction of ion chromatograms and for the quantification of peak areas of heavy Q-peptides and light native peptides.

## 3 Results

### 3.1 Construction of *Chlamydomonas* strains overexpressing sedoheptulose-1,7-bisphosphatase (SBP1)

The *Chlamydomonas SBP1* gene encodes sedoheptulose-1,7-bisphosphatase of the CBC. We chose to use the genomic version of the gene including all seven exons and six introns to adapt it to the MoClo syntax (Weber et al., 2011;Patron et al., 2015). For this, we followed the protocol suggested previously (Schroda, 2019), which required two PCR amplifications to alter sequences around the start and stop codons and to remove an internal BpiI recognition site (Figure 1A). Using the *Chlamydomonas* MoClo toolkit (Crozet et al., 2018), the domesticated *SBP1* gene was equipped with the strong constitutive *HSP70A-RBCS2* fusion promoter (A(Δ-467)-R) and the *RPL23* terminator (Strenkert et al., 2013;Lopez-Paz et al., 2017). We generated two variants, one encoding a 3xHA tag at the C-terminus (SBP1-3xHA), the other lacking any tags (SBP1-mStop) (Figure 1A). HA-tagged proteins are easy to screen for, because the anti-HA antibody used reacts strongly with the 3xHA tag and has little background on immunoblots with *Chlamydomonas* total proteins. This allows assessing the frequency and variance with which transformants express the transgenic protein, and whether it has the expected size. This information can then be used for the screening of transformants expressing the untagged transgenic protein, which is the preferable variant because the 3xHA tag might interfere with the protein’s function. After adding an *aadA* cassette to the constructs (Figure 1A), they were transformed into the *Chlamydomonas* UVM4 strain that expresses transgenes efficiently (Neupert et al., 2009). Of the 12 SBP1-3xHA transformants screened, three did not express the transgene and five expressed it to high levels (Figure 1B). A similar pattern was observed for the 12 SBP1-mStop transformants, of which three appeared not to express the transgene and three expressed it to high levels. The two best-expressing transformants of each construct were selected for further analyses.

**Figure 1.**
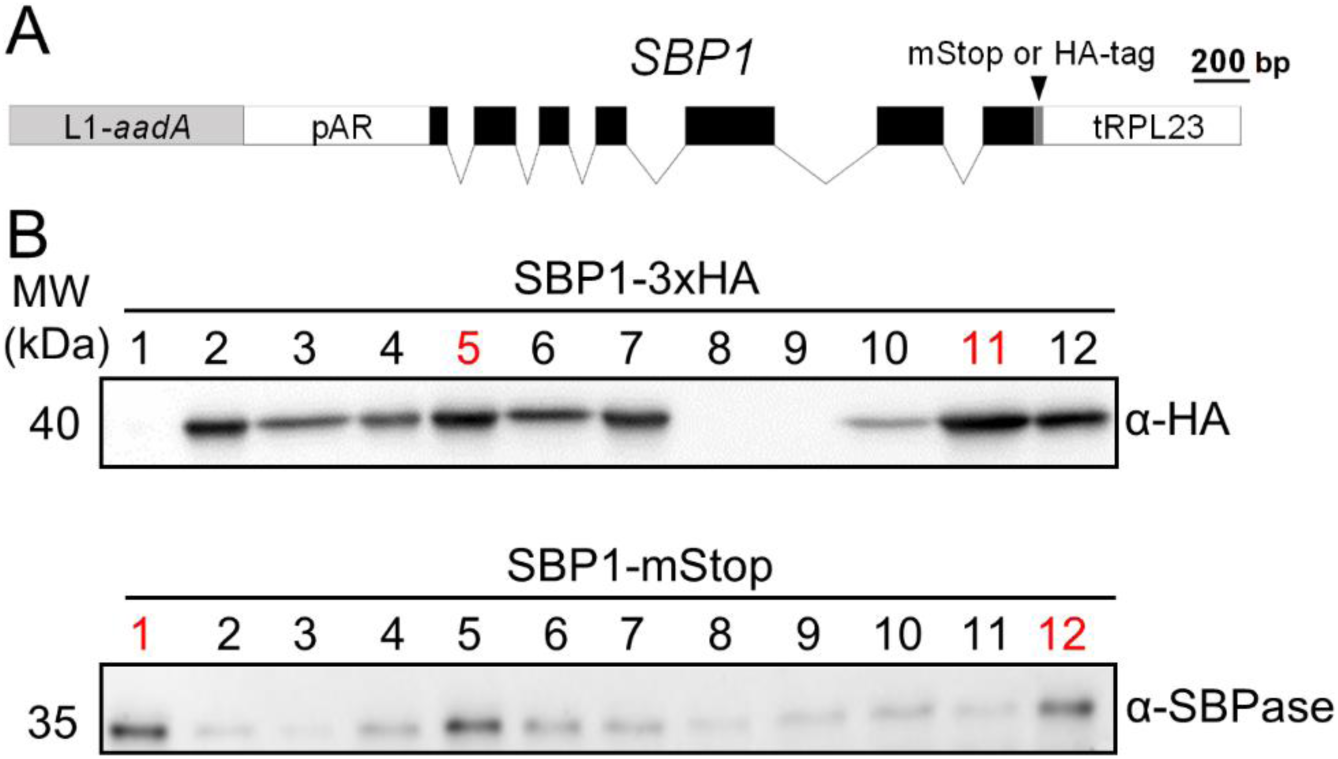
Generation of *Chlamydomonas* lines overexpressing SBP1. **(A)** SBP1 construct used for transformation. The 2172 bp *SBP1* ORF (exons shown as black boxes), interrupted by all native *SBP1* introns (thin lines), was domesticated to generate a level 0 module for the MoClo strategy. Using MoClo, the *SBP1* ORF was equipped with the *HSP70A-RBCS2* promotor and the RPL23 terminator (pAR and tRPL23, respectively, white boxes) and with a 3xHA-tag or without a tag (mStop) (grey box), giving rise to two level 1 constructs. These were combined with another level 1 construct containing the *aadA* gene conferring resistance to spectinomycin (light grey box) to yield the final level 2 constructs for transformation **(B)** Screening of transformants overexpressing SBP1. The UVM4 strain was transformed with the level 2 constructs shown in (A). Total cell proteins from 12 spectinomycin resistant transformants recovered with each construct were extracted and proteins corresponding to 1.5 µg chlorophyll were analyzed by immunoblotting using anti-HA or anti-SBPase antibodies. Transformants exhibiting highest expression levels for SBP1-3HA or SBP1-mStop (red) were used for further analysis.

### 3.2 Monitoring SBP1-overexpressing lines for improved photosynthetic rate and growth

We first tested the four SBP1-overexpressing transformants for improved growth. As elevated SBPase activity has resulted in improved growth particularly under high light and high CO_2_ conditions (Miyagawa et al., 2001;Lefebvre et al., 2005;Tamoi et al., 2006;Ichikawa et al., 2010;Gong et al., 2015;Ogawa et al., 2015;Driever et al., 2017;De Porcellinis et al., 2018), we chose to grow the transformants under mixotrophic conditions with acetate in the medium at a light intensity of 150 µmol photons m^−2^ s^−1^ (our standard growth light intensity is 40 µmol photons m^−2^ s^−1^). When *Chlamydomonas* cells use acetate as a carbon source, they generate a CO_2_-enriched environment by respiration. As shown in Figure 2A, both SBP1-mStop transformants (St1 and St12) accumulated significantly higher (p < 0.001) culture volumes after 44 h and 52 h of growth and therefore reached stationary phase about 14 h earlier than the UVM4 recipient strain. Growth of the HA5 transformant did not differ from that of UVM4 and growth of HA11 even lagged behind that of UVM4. To test whether the enhanced growth rate of the SBP1-mStop transformants was due to an improved photosynthetic rate, we monitored the photosynthetic light response curves for UVM4 and the two SBP1-mStop lines. For this, we measured oxygen evolution as a function of applied light intensity under mixotrophic growth conditions (Figure 2B). Rates of oxygen evolution of the UVM4 strain were comparable with those measured earlier in another strain background (CC-125), and both strains exhibited maximal oxygen evolution at 450 µmol photons m^−2^ s^−1^ (Wykoff et al., 1998). While UVM4 and the four transformants evolved oxygen with similar rates at light intensities of up to 183 µmol photons m^−2^ s^−1^, the SBP1-mStop lines started to evolve more oxygen at light intensities exceeding 183 µmol photons m^−2^ s^−1^ and this became significant (p < 0.05) at light intensities of 520 µmol photons m^−2^ s^−1^ and above. Under photoautotrophic growth conditions at a light intensity of 150 µmol photons m^−2^ s^−1^ we observed no difference in growth between all strains, presumably because they were CO_2_ limited (Figure 2C). We found no differences in chlorophyll content between the strains (t-test, p > 0.05, n=3).

**Figure 2.**
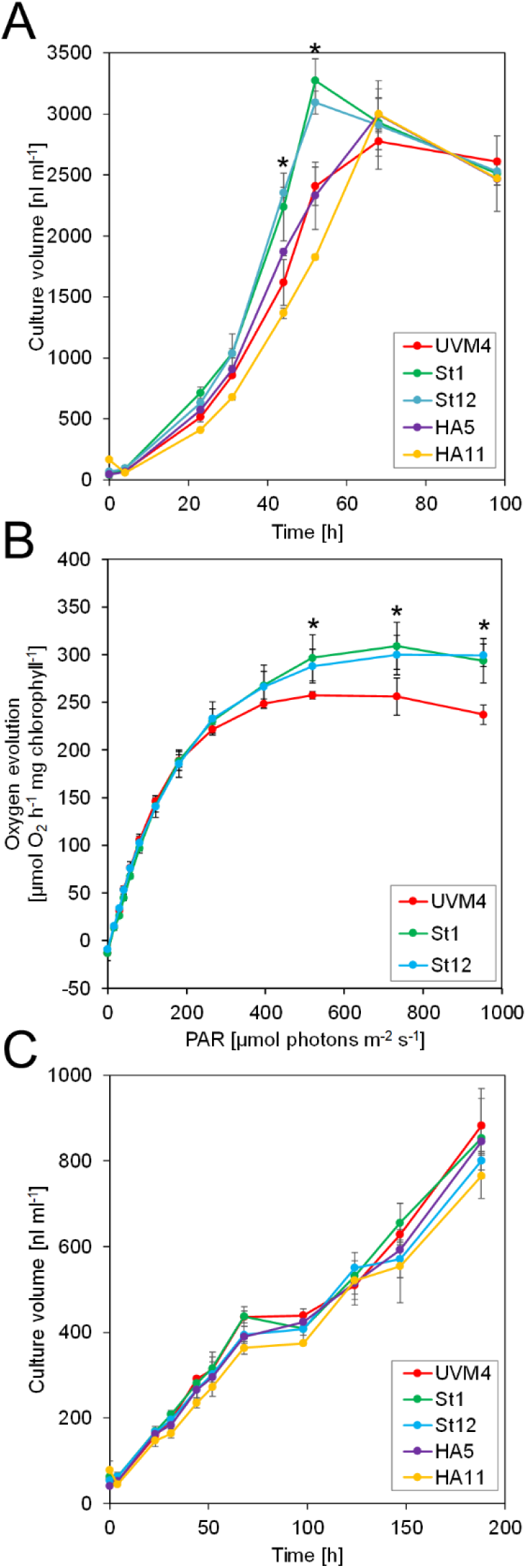
Growth and light response curves of SBP1-overexpressing lines versus the UVM4 recipient strain. **(A)** Growth curves under mixotrophic conditions. Cultures were inoculated in TAP medium at a density of 3 × 10^5^ cells ml^−1^ and incubated on a rotatory shaker for 4 days at a light intensity of 150 µmol photons m^−2^ s^−1^. The culture volume (cell density × cell size) was determined with a Coulter counter. Error bars indicate SD, n = 3. Asterisks indicate significant differences to the UVM4 strain, p < 0.001 (one-way ANOVA with Dunnett’s multiple comparison test). **(B)** Light response curves. Cells were grown mixotrophically to mid-log phase and oxygen evolution at the indicated light intensities was measured on a Mini-PAM II with needle-type oxygen microsensor OXR-50. Error bars indicate SD, n = 3. Asterisks indicate significant differences to the UVM4 strain, p < 0.05 (one-way ANOVA with Dunnett’s multiple comparison test). **(C)** Growth curves under photoautotrophic conditions. Cultures were inoculated in HMP medium at a density of 3 × 10^5^ cells ml^−1^ and incubated on a rotatory shaker for 10 days at a light intensity of 150 µmol photons m^−2^ s^−1^. The culture volume (cell density × cell size) was determined with a Coulter counter. Error bars indicate SD, n = 3.

### 3.3 Absolute quantification of all CBC enzymes in *Chlamydomonas* by the QconCAT strategy

We observed improved growth for the SBP1-mStop transformants but not for the SBP1-3xHA transformants. We reasoned that this could have been due to higher SBP1 expression levels in the former, or due to a negative effect of the 3xHA tag on the protein’s function in the latter. To distinguish between these possibilities and to elucidate whether SBP1 overexpression affected the expression of the other ten CBC enzymes, we quantified the absolute levels of all CBC enzymes in the UVM4 recipient strain and the four SBP1-overexpressing transformants with the QconCAT strategy. With this approach, using a single QconCAT protein (PS-Qprot), we already had determined the absolute cellular quantities of the complexes involved in the photosynthetic light reactions and of the Rubisco rbcL and RBCS subunits (Hammel et al., 2018). We designed a QconCAT protein (CBC-Qprot) that covered each of the missing ten CBC enzymes with two or three proteotypic tryptic Q-peptides (Supplemental Figure 1A; Supplemental Dataset 1). These Q-peptides have been detected by LC-MS/MS in earlier studies with good ion intensities and normal retention times. We had selected them before the d::pPOP algorithm for predicting ionization propensities was available (Zimmer et al., 2018) and therefore some peptides are not the very best choice (see d::pPop ranks and scores in Table 1).

**Table 1.**
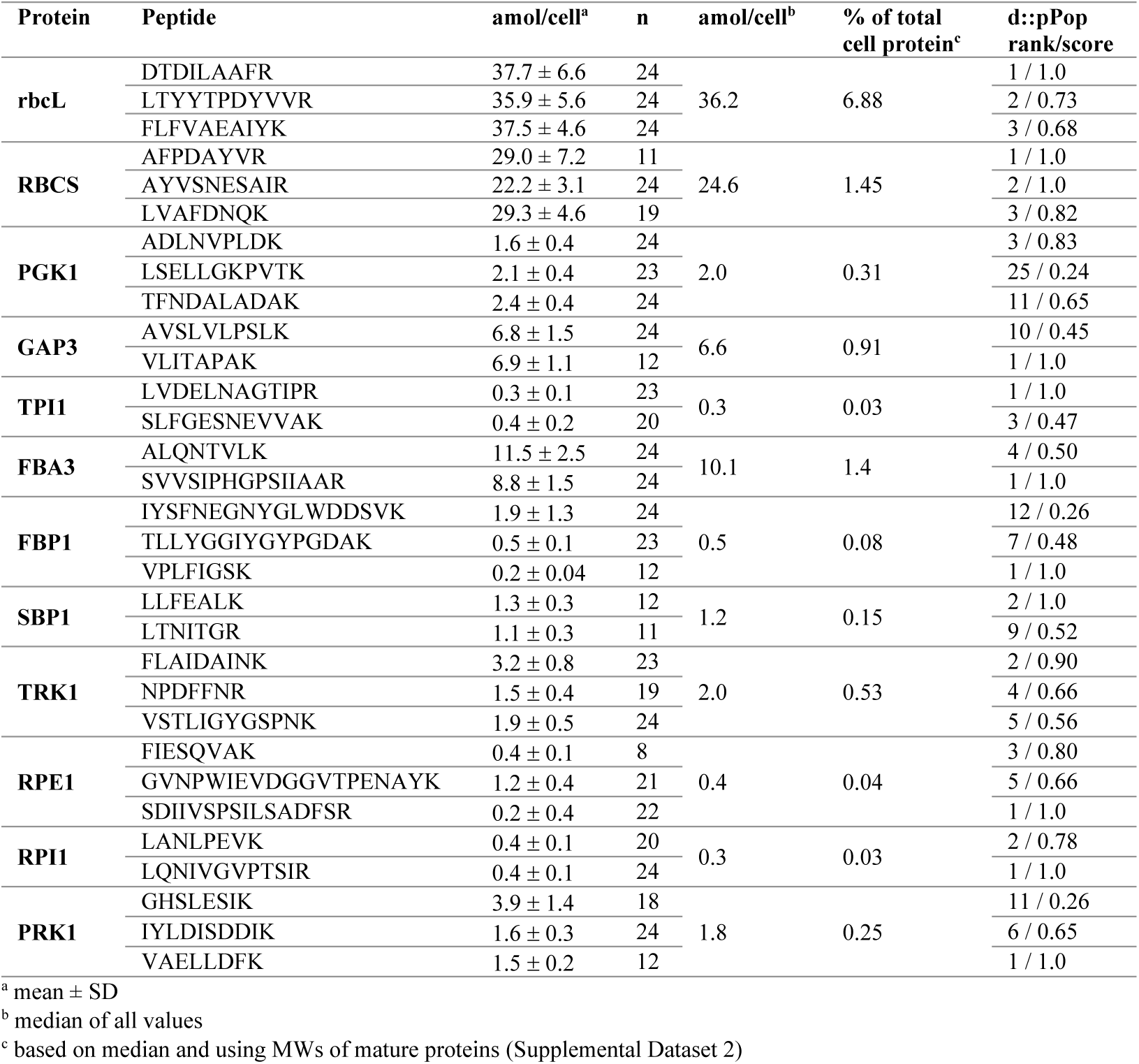
Absolute quantification of Calvin-Benson-Cycle proteins in the *Chlamydomonas* UVM4 strain.

The 47.9-kDa CBC-Qprot was expressed as ^15^N-labeled protein in *E. coli* and purified via the tandem-hexa-histidine tag at its C-terminus. The protein was further purified by preparative electrophoresis on an SDS-polyacrylamide gel, followed by electroelution of the protein from the excised gel band and spectroscopic quantification. Correct quantification and purity was verified by separating the CBC-Qprot together with a BSA standard on an SDS-polyacrylamide gel and staining with Coomassie blue (Supplemental Figure 1B). The CBC-Qprot was then tryptically digested and released peptides analyzed by LC-MS/MS using a short 6-min gradient (Supplemental Figure 1C). The latter shows that the Q-peptides separated with characteristic retention times and ion intensities. Despite the strict 1:1 stoichiometry of the peptides, the areas of the extracted ion chromatograms (XICs) varied by a factor of up to 370.

Four different amounts of the ^15^N-labeled PS-Qprot (Hammel et al., 2018) and the CBC-Qprot were mixed with 20 µg of (^14^N) whole-cell proteins extracted from samples of UVM4 and the four transformants taken 22 h after inoculation in the experiment shown in Figure 1A (early log phase). We employed only one preparation of the QconCAT proteins, but up to six independent samples of *Chlamydomonas* cells. Mixed proteins were precipitated with acetone, followed by tryptic digestion in urea and LC-MS/MS analysis on 45-min analytical gradients. The ion chromatograms of heavy Q-peptide and light native peptide pairs were extracted, XICs quantified, and ratios calculated (Supplemental Dataset 2).

Based on the Q-peptide to native peptide ratios and the known amounts of spiked-in QconCAT proteins, the abundances of the native peptides in the sample were calculated (in femtomoles per µg cell protein) (Supplemental Dataset 2). We determined that a *Chlamydomonas* UVM4 cell contains 27.6 ± 1.7 pg protein (SD, n = 6), which allowed us to calculate the absolute amount of each peptide in attomol per cell (Table 1). We used the median of all quantification values of the 2-3 Q-peptides per protein (23 to 72 values) to get an estimate for the abundance of each CBC protein per cell (Table 1). Moreover, based on these median values and the molecular weight of the mature proteins, the fraction of each target protein in the whole-cell protein extract was estimated (Table 1), revealing that CBC enzymes represent ∼11.9% of total cell protein in *Chlamydomonas* (Supplemental Dataset 2). This procedure was repeated for all four SBP1-overexpressing transformants and the log2-fold change of the abundance of each CBC enzyme in the transformants versus the UVM4 strain was calculated (Figure 3; Supplemental Dataset 2). It turned out that SBP1 was significantly (p < 0.01) overexpressed in all transformants (HA5: 1.6-fold; HA11: 1.7-fold; St1: 3-fold; St12: 2.2-fold). Except for the Rubisco subunits, levels of all other CBC enzymes were not significantly different between the SBP1-overexpressing transformants and the UVM4 strain. Compared to UVM4, transformant HA11 had significantly lower RBCS levels (but only by 8%), and transformant St12 had significantly lower levels of RBCS and rbcL (by about 40%).

**Figure 3.**
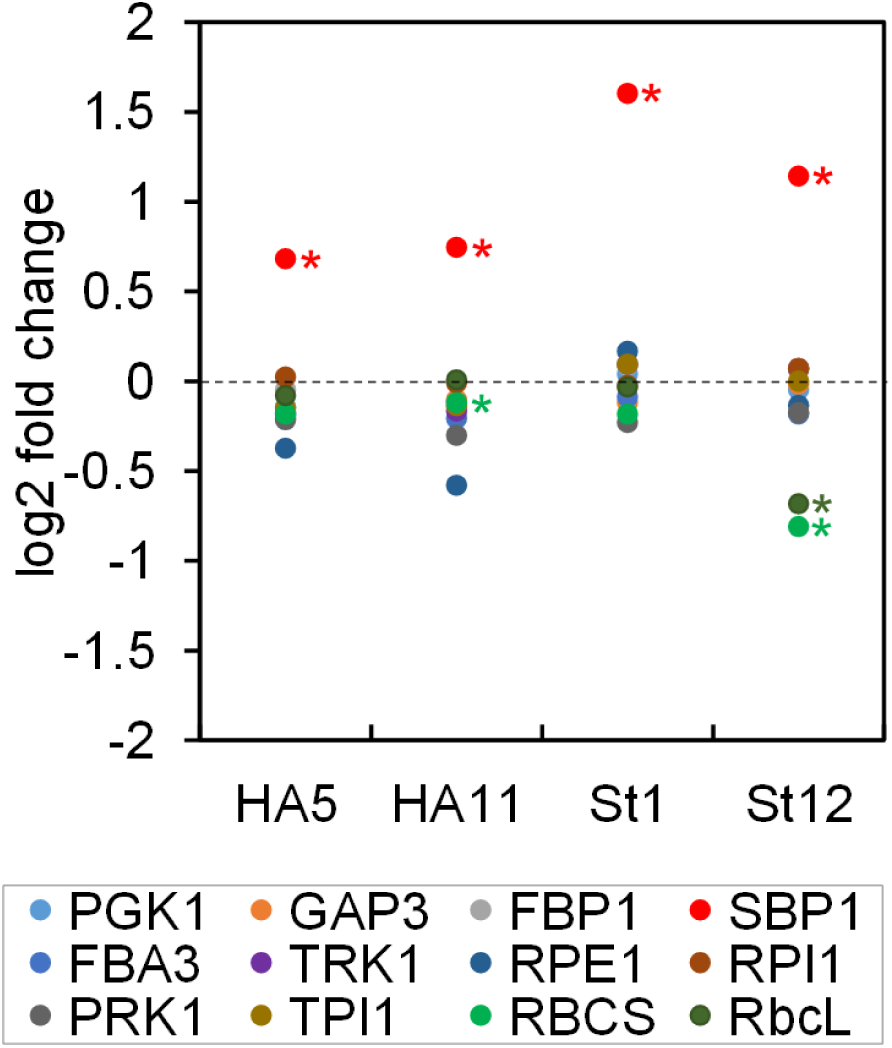
Changes in abundance of CBC enzymes in SBP1-overexpressing lines versus the UVM4 recipient strain. The abundance of the CBC enzymes in transformants HA5 and HA11, generated with the SBP1-3xHA construct, and in transformants St1 and St12, generated with the SBP1-mStop construct, was determined using the QconCAT strategy. Abundances relative to those in the UVM4 recipient strain were log2 transformed and plotted. Asterisks designate significant differences between enzymes in the transformants versus the UVM4 strain (Kruskal-Wallis test with Dunn’s post hoc test, p < 0.01). PGK, phosphoglycerate kinase; GAP, glyceraldehyde-3-phosphate dehydrogenase; TPI, triose phosphate isomerase; FBA/SBA, fructose-1,6-bisphosphate aldolase/sedoheptulose-1,7-bisphosphate aldolase; FBP, fructose-1,6-bisphosphatase; TRK, transketolase; SBP, sedoheptulose-1,7-bisphosphatase; RPE, ribulose-5-phosphate 3-epimerase; RPI, ribose-5-phosphate isomerase; PRK, phosphoribulokinase; rbcL, ribulose bisphosphate carboxylase/oxygenase large subunit; RBCS, ribulose bisphosphate carboxylase/oxygenase small subunit.

### 3.4 Estimation of substrate binding sites per CBC enzyme

In a previous study, we had determined the levels of all CBC metabolites in *Chlamydomonas* cells during an increase in light intensity within the range where irradiance remains limiting for photosynthesis (Mettler et al., 2014). In that study, also the concentrations of the CBC enzymes in the chloroplast were estimated based on the empirical protein abundance index (emPAI) (Ishihama et al., 2005). These data sets allowed estimating the number of substrate binding sites per CBC enzyme 20 min after increasing the light intensity, when flux through the cycle was maximal (Mettler et al., 2014). To compare the emPAI-derived data with the QconCAT-derived data, we calculated the concentration of each CBC enzyme in the chloroplast based on the absolute quantities determined here and the assumption that a *Chlamydomonas* cell has a volume of 270 µm^3^, of which about half is occupied by the chloroplast (Weiss et al., 2000) (Table 2). While the concentration of rbcL determined by Mettler et al. (2014) matched that determined here with the QconCAT approach very well, the concentrations of all other CBC enzymes were strongly overestimated (between 8.8-fold for FBA3 and 34.5-fold for PRK1) (Table 2). To re-estimate the number of substrate binding sites per CBC enzyme, we used the concentrations of the CBC enzymes in the chloroplast determined here and the CBC metabolite data determined earlier 20 min after the light shift to 145 µmol photons m^−2^ s^−1^ (Mettler et al., 2014) (Table 2). Although the growth conditions differed slightly between the two studies, metabolite levels do not vary greatly in *Chlamydomonas* in this irradiance range (Mettler et al., 2014, Supplemental Figure 12).

**Table 2.**
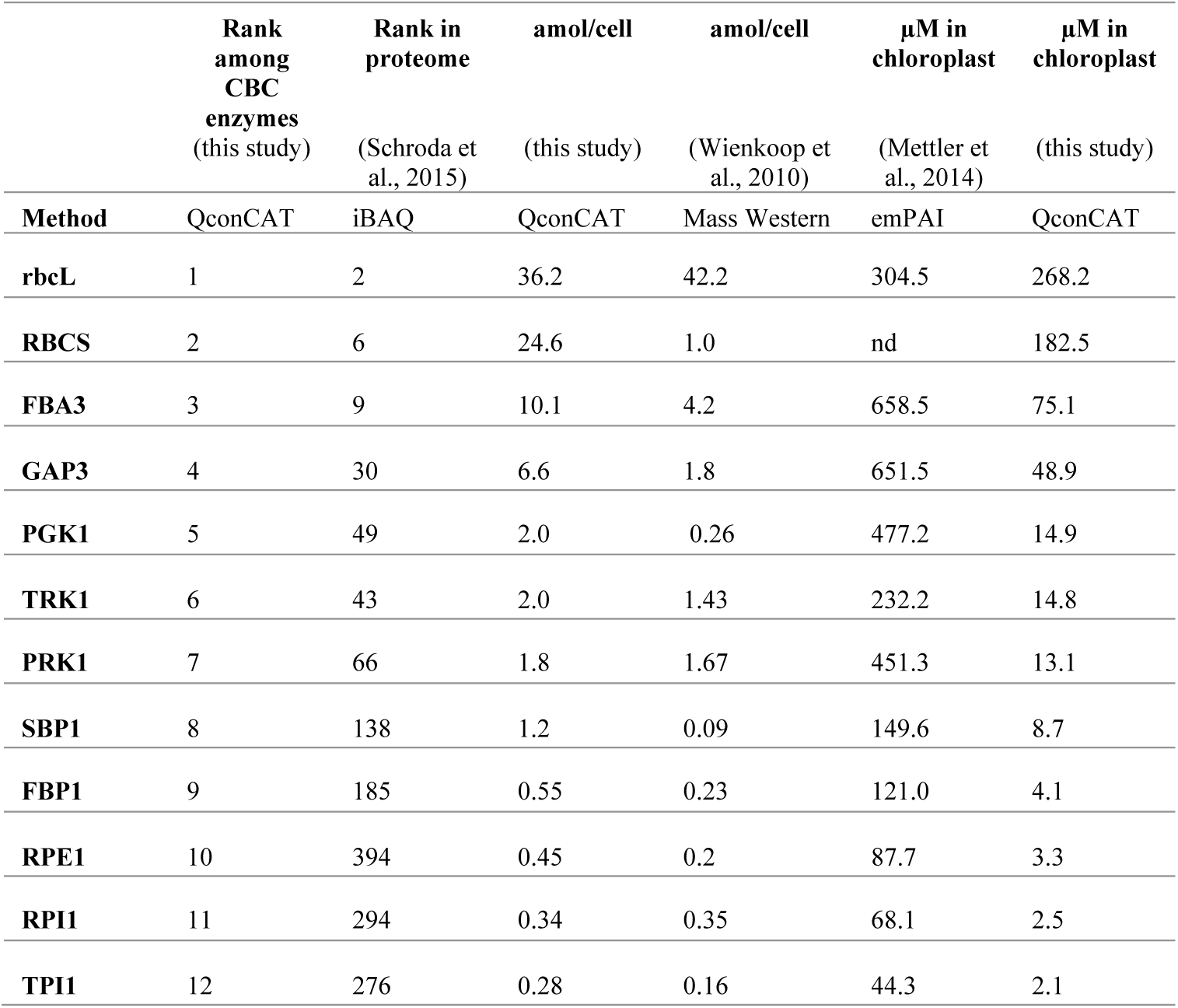
Comparison of CBC enzyme abundances and concentrations in *Chlamydomonas* determined in different studies by different methods.

This re-analysis revealed that some CBC intermediates are indeed present at lower concentrations than the estimated binding site concentration of the enzymes for which they act as substrates (1,3-bisphosphoglycerate (BPGA) compared to PGK1 and GAP3; glyceraldehyde 3-phosphate (GAP) and erythrose 4-phosphate (E4P) compared to FBA3), some are at only slightly (<4.5-fold) higher concentrations than the respective binding site (GAP and E4P compared to TRK; ribulose-5-phosphate (Ru5P) compared to PRK1; fructose-1,6-bisphosphate (FBP) compared to FBA3). However, most of the other CBC intermediates are present at considerably higher concentrations than the respective estimated binding site concentration (Figure 4).

**Figure 4.**
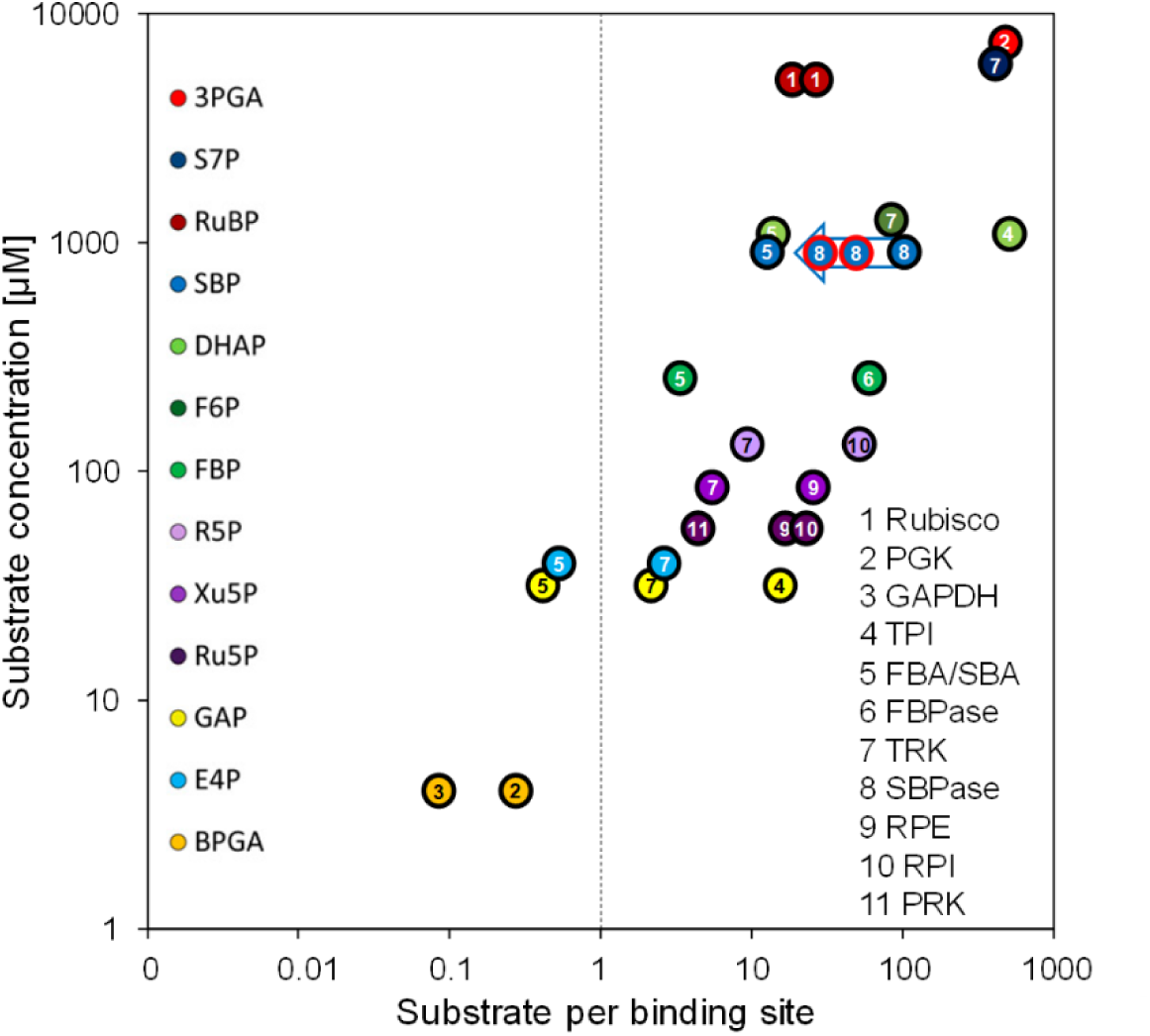
Substrate per binding sites versus substrate concentrations. Substrates of CBC reactions, determined by LC-MS/MS, were taken from Mettler et al. (2014). Binding sites of CBC enzymes were calculated based on the QconCAT data (Table 2). Substrate per binding site values of CBC reactions are plotted against the substrate level 20 min after the light intensity was increased from 41 to 145 µmol photons m^−2^ s^−1^. For enzymes that catalyze readily reversible reactions (TPI, FBA/SBA, TRK, RPI, RPE) the relation to product level is also shown. To facilitate comparison, the same numbering and color code as used in Mettler et al. (2014) was adopted. The blue arrow shows the estimates for substrates per binding site for the SBP1 overexpressing lines St1 and St12 (encircled in red). The two values for Rubisco are based on the slightly different quantification values for the large and small subunits.

## 4 Discussion

### 4.1 The modularity of the MoClo approach and the use of *Chlamydomonas* as a model facilitate the iterative process of genetic engineering towards improving plant productivity

Here we present a workflow for rapid and efficient metabolic engineering towards improving plant biomass production, with the overexpression of native *Chlamydomonas* SBPase (SBP1) in *Chlamydomonas* as a proof-of-concept. We used the Modular Cloning (MoClo) strategy for construct generation and employed the part library established recently (Weber et al., 2011;Crozet et al., 2018). The one-step, modular assembly of multiple genetic parts allowed generating complex constructs rapidly and with variations: one coding for SBP1 with a 3xHA tag and one lacking any tags. This double strategy was well chosen, as the variant with a C-terminal 3xHA tag did not result in enhanced photosynthetic rates and biomass production, while the variant lacking a tag did (Figures 2A and 2B). In the two transformant lines tested for each construct, tagged SBP1 was overexpressed 1.6 to 1.7-fold while the untagged form was overexpressed ∼2.2- and ∼3-fold (Figure 3). Therefore, it is possible that in *Chlamydomonas* SBPase must be expressed to levels higher than 1.7-fold to improve the photosynthetic rate. Alternatively, the C-terminal 3xHA tag interfered with SBP1 function. We favor the latter explanation, because SBPase overexpression giving rise to at most 2-fold increased activities already had positive effects on photosynthetic rates and biomass accumulation in tobacco (Lefebvre et al., 2005;Tamoi et al., 2006;Rosenthal et al., 2011), *Dunaliella bardawil* (Fang et al., 2012), and wheat (Driever et al., 2017). Like for these models, increased photosynthetic rates and biomass accumulation were observed in *Chlamydomonas* lines overexpressing SBP1 only if cells were grown at higher light intensities (150 µmol photons m^−2^ s^−1^) and elevated CO_2_ concentrations (in the presence of acetate) (Figure 2). Hence, under these conditions SBPase levels represent a bottleneck in flux through the CBC in *Chlamydomonas* as in the other plant models. Therefore, the results obtained with *Chlamydomonas* readily apply to other alga and land plants.

SBP1 represents 0.15% of total cell protein in *Chlamydomonas* (Table 1), i.e. the transgenic protein in the best SBP1-overexpressing line makes up 0.3% of total cell protein. It is likely that the screening of more transformants would have allowed recovering lines with even higher expression levels. Furthermore, a *SBP1* gene re-synthesized with optimal codon usage and the three *RBCS2* introns probably would have allowed higher expression levels (Barahimipour et al., 2015;Schroda, 2019). By combining the MoClo strategy with *Chlamydomonas* as a model, a complete cycle of the iterative process of construct design and assembly, transformation, screening, and phenotype test can be achieved in as little as 6 weeks.

### 4.2 QconCAT-based quantitative proteomics allows monitoring effects of SBP1 overexpression on the accumulation of other CBC enzymes

Increased activities of SBPase by overexpressing SBPase alone or BiBPase from cyanobacteria had no effect on the levels or activities of selected other CBC enzymes in tobacco (Miyagawa et al., 2001;Lefebvre et al., 2005;Rosenthal et al., 2011), lettuce (Ichikawa et al., 2010) or wheat (Driever et al., 2017). In contrast, overexpression of BiBPase from *Synechocystis* in *Synechococcus* resulted in increased activities of Rubisco (2.4-fold) and aldolase (1.6-fold) as well as increased protein levels of rbcL (∼3-fold), TPI (1.5-fold) and RPI (1.4-fold) (De Porcellinis et al., 2018). Similarly, overexpressing SBPase leading to up to 1.85-fold higher SBPase activity in *Arabidopsis* resulted in elevated FBPA activity and protein levels (Simkin et al., 2017a). We employed the QconCAT approach to determine absolute quantities of all other ten CBC enzymes and found no consistent changes between wild type and the four SBP1-overexpressing lines (Figure 3). Only the St12 line with ∼2.2-fold higher SBP1 expression had a significant ∼40% reduction of both Rubisco subunits rbcL and RBCS. Since both Rubisco subunits were unaffected in line St1 with ∼3-fold higher SBP1 levels, SBP1 overexpression cannot be the cause for the reduced accumulation of Rubisco in line St12. More likely, a gene required for Rubisco expression, assembly or stability was destroyed by the integration of the SBP1 expression vector. It is surprising that photosynthetic rate and biomass accumulation was increased to a similar extent in lines St1 and St12 despite the reduced Rubisco levels in line St12 (Figure 2). This indicates that Rubisco levels are not limiting CBC flux in *Chlamydomonas*, in line with previous observations in *Chlamydomonas* that reducing Rubisco to almost 50% of wild-type levels enabled full photosynthetic growth (Johnson, 2011).

### 4.3 CBC enzymes exhibit a larger abundance range than estimated earlier

In addition to looking at possible effects of SBP1-overexpression on the expression of other CBC enzymes, the QconCAT approach allowed for the quantification of absolute levels of CBC enzymes in *Chlamydomonas* cells. With this strategy, we had already determined absolute quantities of rbcL and RBCS in another cell wall-deficient strain background (CC-1883) (Hammel et al., 2018). There, absolute amounts of rbcL and RBCS were ∼1.4-fold lower than in UVM4 cells. However, CC-1883 cells also had a ∼1.3-fold lower protein content than UVM4 cells, such that the fraction of rbcL and RBCS in total cell protein are about comparable (6.6% and 1.3% in CC-1883 versus 6.88% and 1.45% in UVM4, respectively).

The abundance of all CBC enzymes in *Chlamydomonas* cells has been estimated earlier. One study used “Mass Western”, which is based on the spiking-in of known amounts of heavy isotope-labeled Q-peptides into tryptic digests of whole-cell proteins followed by LC-MS/MS analysis (Wienkoop et al., 2010). The other studies used the emPAI (empirical protein abundance index) and iBAQ (intensity-based absolute quantification) approaches on quantitative shotgun proteomics datasets (Mettler et al., 2014;Schroda et al., 2015).

The iBAQ-based ranking of protein abundances exactly reflects the quantities of the more abundant CBC enzymes determined here by the QconCAT approach (Table 2). Only the low-abundance CBC enzymes RPE1, RPI1 and TPI1 were ranked by iBAQ in the opposite order of their abundance determined by the QconCAT method. Most likely, this is due to the impaired accuracy of the iBAQ approach for less-abundant proteins (Soufi et al., 2015).

The absolute quantities determined by Mass Western roughly matched those determined with the QconCAT approach, except for RBCS, PGK1 and SBP1, which were 24.6-fold, 7.7-fold, and 13.3-fold lower (Table 2). As suggested earlier (Hammel et al., 2018), this discrepancy can be explained by an incomplete extraction of some proteins from whole-cell homogenates with the extraction protocol employed (Wienkoop et al., 2010).

In the study by Mettler et al. (2014), the cellular abundance of rbcL was estimated by densitometry on Coomassie-stained SDS-gels and was used to normalize the emPAI-derived quantification values of the other CBC enzymes. The estimated abundance of rbcL matches that determined here via the QconCAT approach (Table 2). However, the emPAI-derived values for the other CBC enzymes are much higher than those determined by QconCAT (up to 34.5-fold higher for PRK1). A likely cause for this strong overestimation is that proteins of very high abundance tend to exhibit a saturated emPAI signal (Ishihama et al., 2008). Consequently, the range of concentrations of CBC enzymes is much larger than estimated earlier. For example, the difference between the most abundant CBC protein rbcL and the least abundant TPI1 is 128-fold rather than only 7-fold (Table 2). The strong overestimation of many CBC enzymes by the emPAI approach challenges the conclusion that many CBC intermediates are present at concentrations that are far lower than the estimated binding site concentration of the enzymes for which they act as substrates (Mettler et al., 2014). For example, the concentration of sedoheptulose-1,7-bisphosphate (SBP) is ∼106-fold higher than that of SBP1 rather than only ∼6-fold as estimated previously (Figure 4). Moreover, comparisons of the *in vivo* SBP concentration and the modelled *in vivo* K_m_ for SBP1 indicate that SBP1 is likely to be near-saturated *in vivo* (Mettler et al., 2014). Hence, flux at SBPase is likely restricted by the degree of post-translational activation of SBP1 and SBP1 abundance. This explains better why an increase in CBC flux can be achieved by increasing SBP1 protein concentrations.

Still correct is that the concentration of GAP is below or slightly above the concentration of substrate binding sites of FBA3 and TRK1 (0.4-fold and 2.2-fold, respectively), as is E4P compared to FBA3 and TRK1 (0.5-fold and 2.7-fold, respectively). Furthermore, ribulose-5-phosphate (Ru5P) is only 4.3-fold above the binding site concentration of PRK, indicating that increased flux in the regeneration phase of the CBC to increase Ru5P levels will aid increase RuBP formation and fixation of CO_2_. The low concentration of these key CBC intermediates relative to their enzyme binding sites, together with the low concentration of these and further CBC intermediates relative to the likely *in vivo* K_m_ values of CBC enzymes (Mettler et al., 2014), explains how RuBP regeneration speeds up when rising light intensity drives faster conversion of 3-phosphoglycerate to GAP.

### 4.4 Outlook

The next step would be to stack multiple transgenes for the overexpression of CBC enzymes that in SBP1-overexpressing lines potentially become new bottlenecks for flux through the cycle. Indicative for this scenario is the finding that SBPase overexpression in *Arabidopsis* entailed an overexpression of FBPA (Simkin et al., 2017a). Moreover, overexpression of BiBPase in *Synechococcus* came along with an increase in levels of RPI and TPI (De Porcellinis et al., 2018), which are the CBC enzymes of lowest abundance in *Chlamydomonas* (Tables 1 and 2). More candidates for multigene stacking might be PGK1, TRK1, TPI1 and FBP1, whose substrates are in largest excess of the substrate binding sites (Figure 4). To our knowledge, there are yet no reports on the overexpression of PGK, RPI, and TPI (Simkin et al., 2019). Two studies report no or even negative effects upon TRK overexpression in rice and tobacco, respectively (Khozaei et al., 2015;Suzuki et al., 2017). Positive effects of FBPase overexpression on photosynthetic rates and biomass accumulation were reported for numerous plant models – except for *Chlamydomonas* where FBP1 overexpression in the chloroplast had negative effects (Dejtisakdi and Miller, 2016). Apparently, the highly complex regulation of the CBC and its central role in cellular metabolism make predictions difficult. This is highlighted by recent work, indicating that the balance between different steps in the CBC varies from species to species (Arrivault et al., 2019;Borghi et al., 2019). Therefore, experimental test is the route of choice that with the combination of MoClo and *Chlamydomonas* can be pursued easily.

## 6 Author Contributions

A.H. performed all experiments. F.S. designed the QconCAT protein and performed the LC-MS/MS analyses. A.H., and D.Z. analysed the data and were supervised by T.M., M.S., and M.Sc. M.Sc. conceived and supervised the work. M.Sc. wrote the article with contributions from all other authors.

## 8 Acknowledgments

We are grateful to Karin Gries for technical help.

---

**Supplementary Figure 1.**
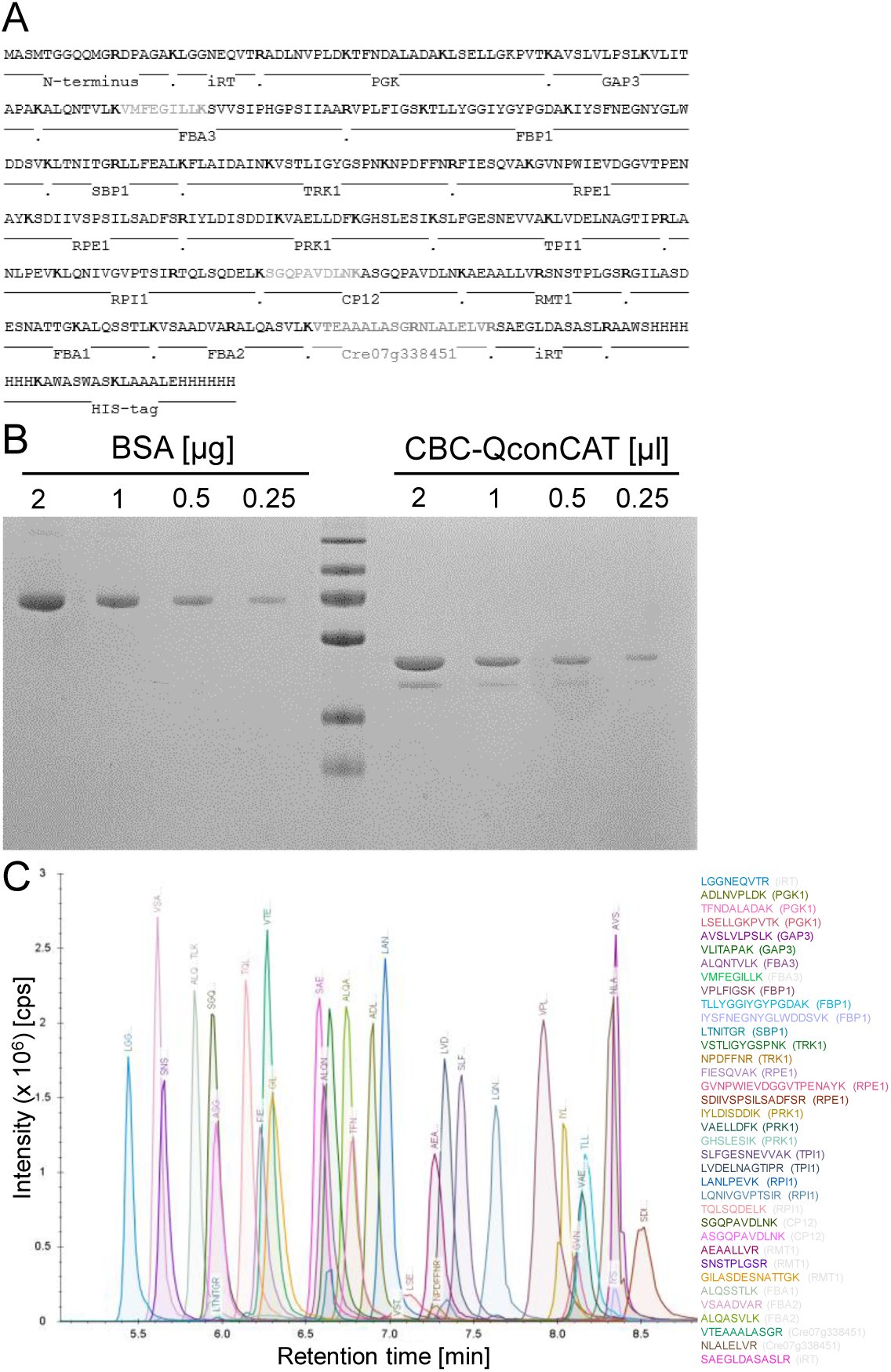
Design and production of the Calvin-Benson-Cycle (CBC) QconCAT protein. **(A)** Sequence of the CBC-Qprot and protein source of selected Q-peptides. Peptides in grey were erroneously included (CP12), have a proline following the tryptic cleavage site in the native context (FBA3), or belong to a fructose-bisphosphatase not involved in the Calvin-Benson-Cycle (Cre07.g338451). iRT peptides can be used for retention time aligment. **(B)** The purified, ^15^N-labeled CBC-Qprot was quantified on a NanoDrop spectrophotometer and the concentration adjusted to 1 μg/μl. The indicated volumes of the CBC-Qprot were then separated next to a BSA standard on a 12%-SDS polyacrylamide gel and stained with Coomassie blue. The labeling efficiency of the CBC-Qprot was 99.39% ± 0.37%. **(C)** Extracted ion chromatograms (XICs) of the proteotypic ^15^N labeled Q-peptides derived from the PS-Qprot. The purified protein was tryptically digested and run on a short 6-min HPLC gradient. XICs of the resolved peptides were extracted using the PeakView software (ABSciex). Peptides for which the corresponding protein name is given in grey were not used for quantification. Note that due to the very short not all peptides were detected within the retention time window.

